# Nucleolar Cdc14 Splitting Reflects Recombination Context and Meiotic Chromosome Dynamics

**DOI:** 10.1101/2025.11.27.691012

**Authors:** Patricia Rodríguez-Jiménez, Paula Alonso-Ramos, Isabel Acosta, David Álvarez-Melo, Jesús A. Carballo

**Affiliations:** Institute of Functional Biology and Genomics (IBFG), Spanish National Research Council (CSIC) and University of Salamanca, 37007 Salamanca, Spain; Centro de Biotecnología y Genómica de Plantas, CBGP, CSIC-INIA, Campus de Montegancedo, 28223 Madrid, Spain; Louvain Institute of Biomolecular Science and Technology, Université catholique de Louvain, 1348 Louvain-La-Neuve, Belgium

## Abstract

Chromosome dynamics, recombination, and nucleolar organization intersect during meiotic prophase I, yet how recombination context influences nucleolar architecture remains unclear. We analyzed the nucleolar pool of Cdc14 in *Saccharomyces cerevisiae* under matched prophase-I gating. In a recombination-competent reference (*ndt80Δ*), Cdc14–mCherry formed a predominant single focus with occasional, reversible two-focus episodes that Nop56–GFP placed within the nucleolar compartment (“nucleolar splitting”). Splitting rose sharply in *dmc1Δ ndt80Δ* and remained high in *spo11–y135f dmc1Δ ndt80Δ*, demonstrating that elevated DSB formation is not required and pointing instead to homolog-engagement state as the key variable. Population checkpoint readouts did not map onto the phenotype: Hop1 phosphorylation was strong in *dmc1Δ*, weak/transient in *ndt80Δ*, and undetectable in *spo11–y135f*, yet splitting was high in *dmc1Δ and spo11–y135f* and low in *ndt80Δ*. Persistence metrics showed that events concentrated early and waned in *ndt80Δ*, whereas *dmc1Δ* and *spo11* backgrounds retained activity into later windows, consistently across thresholds. We propose that nucleolar splitting reflects rheological responses of a nucleolar condensate to chromosome-scale forces that vary with homolog engagement. This view is consistent with DSB-independent contributions from telomere clustering into the bouquet configuration, telomere-led rapid prophase movements, and centromere coupling/pairing as routes that shape force transmission to the rDNA territory. In this sense, the nucleolus emerges as a mesoscale, mechanically sensitive readout of meiotic chromosome dynamics.

## 1. Introduction

Meiosis establishes haploidy through a coordinated prophase I in which homologous chromosomes approach one another, assemble synaptonemal complex (SC), and exchange DNA. Double-strand breaks (DSBs) introduced by Spo11 initiate interhomolog recombination, yet only a subset of intermediates mature into crossovers that obey assurance and interference, implying long-range communication along chromosome length [1–3].

A convergent view is that meiotic patterning emerges from mesoscale condensates assembled on chromosome axes. Pro-crossover assemblies can behave as active droplets whose number and spacing arise through coarsening and growth-competition dynamics, providing a physical basis for interference and assurance; quantitative imaging and theory support diffusion-and-aggregation along synapsed chromosomes as the driver of these emergent patterns [4]. In parallel, conserved regulators diffuse within the SC at rates compatible with signal propagation over chromosomal distances, and wetting-like assembly of the SC central region can organize axis-associated compartments and their mechanical couplings [5–8].

At the break-formation step, the Rec114–Mei4–Mer2 (RMM) module forms DNA-bound condensates with liquid-like features; tuning their material properties modulates Spo11-dependent DSB output, placing phase separation at the heart of meiotic DSB biogenesis [9]. Within this framework, processed DSBs generated in these assemblies recruit the ATR orthologue Mec1, which phosphorylates the axis protein Hop1 to trigger the recombination checkpoint and modifies Zip1 at centromeres to dismantle non-homologous centromere coupling [10–12]. The meiosis-specific recombinase Dmc1 then acts on resected Spo11-dependent breaks to promote interhomolog repair, linking DSB formation to both checkpoint signalling and engagement between homologues [13, 14]. These condensates, assembled on chromosomal cores, therefore reshape the mechanical, diffusive, and signalling landscape in which chromosome pairing and recombination unfold.

Chromosome motion is an active contributor. Telomere-led rapid prophase movements (RPM) and transient telomere clustering into the bouquet configuration transmit cytoskeletal forces to chromosome ends through the LINC axis (Ndj1–Mps3–Csm4 and associated bridges), combining nuclear rotation with directed telomere trajectories to remodel chromosome territories [15–17]. Early in prophase I, Zip1-dependent non-homologous centromere coupling and subsequent homolog-centric arrangements provide alternative attachment topologies that can either facilitate or hinder productive pairing; SC components can persist at centromeres after arm disassembly in some contexts [18]. These DSB-independent features—bouquet, RPM and centromere coupling/pairing—set attachment geometry and driving forces for whole-chromosome motion, thereby shaping the nuclear mechanical environment in which recombination proceeds [12, 15, 18–20].

The nucleolus is a multilayered biomolecular condensate. In budding yeast, the rDNA polymer together with a Cdc14-enriched chromatin domain defines a polymer–polymer phase-separated (PPPS) body that coexists with liquid–liquid phase-separated (LLPS) ribonucleoprotein layers formed by box C/D snoRNP components such as Nop56, which mark the bulk nucleolus; these subcompartments differ in rheology and coarsening behaviour and re-establish architecture rapidly after deformation [21–24]. In this framework, the nucleolus is expected to be sensitive to chromosome-borne forces and boundary conditions. Moreover, because the rDNA array on chromosome XII does not assemble canonical SC, forces routed through telomeres and centromeres can act on the nucleolar territory without SC reinforcement, making nucleolar organization a mesoscale reporter of the prophase-I mechanical landscape [18, 25–28].

Cdc14 is a conserved master cell-cycle phosphatase that undergoes regulated nucleolar sequestration and release to drive late mitotic and meiotic transitions. In budding yeast, nucleolar retention through the RENT complex and stage-specific release waves promote mitotic exit and accurate chromosome segregation [29–35]. Genetic analyses, including work from this laboratory, linked Cdc14 activity to the processing of meiotic recombination intermediates and to the fidelity of meiotic chromosome segregation, suggesting that spatial control of the rDNA-associated pool is functionally coupled to meiotic recombination dynamics [24, 36]. This connection motivates the converse question pursued here: does the recombination/engagement context imprint on the nucleolar organization of the Cdc14 pool during prophase I, and can a defined, reversible redistribution within the nucleolus report the balance between DSB-dependent programmes and DSB-independent drivers of pairing/synapsis, as well as its relation to population-level checkpoint readouts?

Here we confront a central gap. Modern work has established that meiotic recombination is organized by condensates that coarsen along the SC, that DSB output is modulated by condensate material properties, and that whole-chromosome motion is actively driven through telomere-clustering into the bouquet configuration, telomere-led RPM and centromere-based attachments. Yet a direct link between these programmes and the morphodynamics of the largest nuclear condensate, the nucleolus, under matched prophase-I conditions is missing. We therefore ask whether a defined, reversible redistribution within the nucleolus can operate as a mechanically sensitive readout of the pairing/engagement landscape, and how such a readout relates to absolute DSB burden and to bulk checkpoint readouts, and whether it can be established rigorously with a uniform operational definition and an independent compartment marker. Resolving this point would bridge molecular regulation, mesoscale mechanics, and nuclear organization in prophase I, turning a longstanding descriptive gap into a tractable, quantitative question.

## 2. Results

### 2.1. A Recombination-Competent Prophase I Reference Uncovers Nucleolar Cdc14 Dynamics

Building on our previous findings [36], which linked Cdc14 to meiotic recombination, here we examined the converse question: whether the recombination context shapes the nucleolar distribution of Cdc14 during prophase I. To establish a recombination-competent reference anchored in prophase I, we analysed *ndt80Δ* and restricted scoring to prophase-I cells identified by Rec8–GFP, a meiosis-specific kleisin subunit of the cohesin complex that marks chromosome axes in prophase I [37–39]. In this reference background, Cdc14–mCherry accumulated as a bright focus within the nucleolar region (Fig. 1A). Interestingly, in a subset of cells, the nucleolar signal transiently resolved into two spatially separated foci within the same territory and later re-joined within the same series, a dynamic behaviour that we scored as two-focus episodes within the nucleolar territory (Fig. 1B, Supplementary Movie S1), using predefined criteria that required a clear intensity valley between two nucleolar Cdc14–mCherry maxima and a measurable spatial separation between their centroids in individual 15-min frames, scored within a ≤12-h window to limit cumulative illumination (Methods; Supplementary Table S2). A compact summary of the number of prophase I cells scored under these criteria in the *ndt80Δ* reference is provided in Fig. 1C. Thus, in a recombination-competent prophase I background, Cdc14 concentrated as a predominant single nucleolar focus with occasional two-focus episodes.

**Figure 1.**
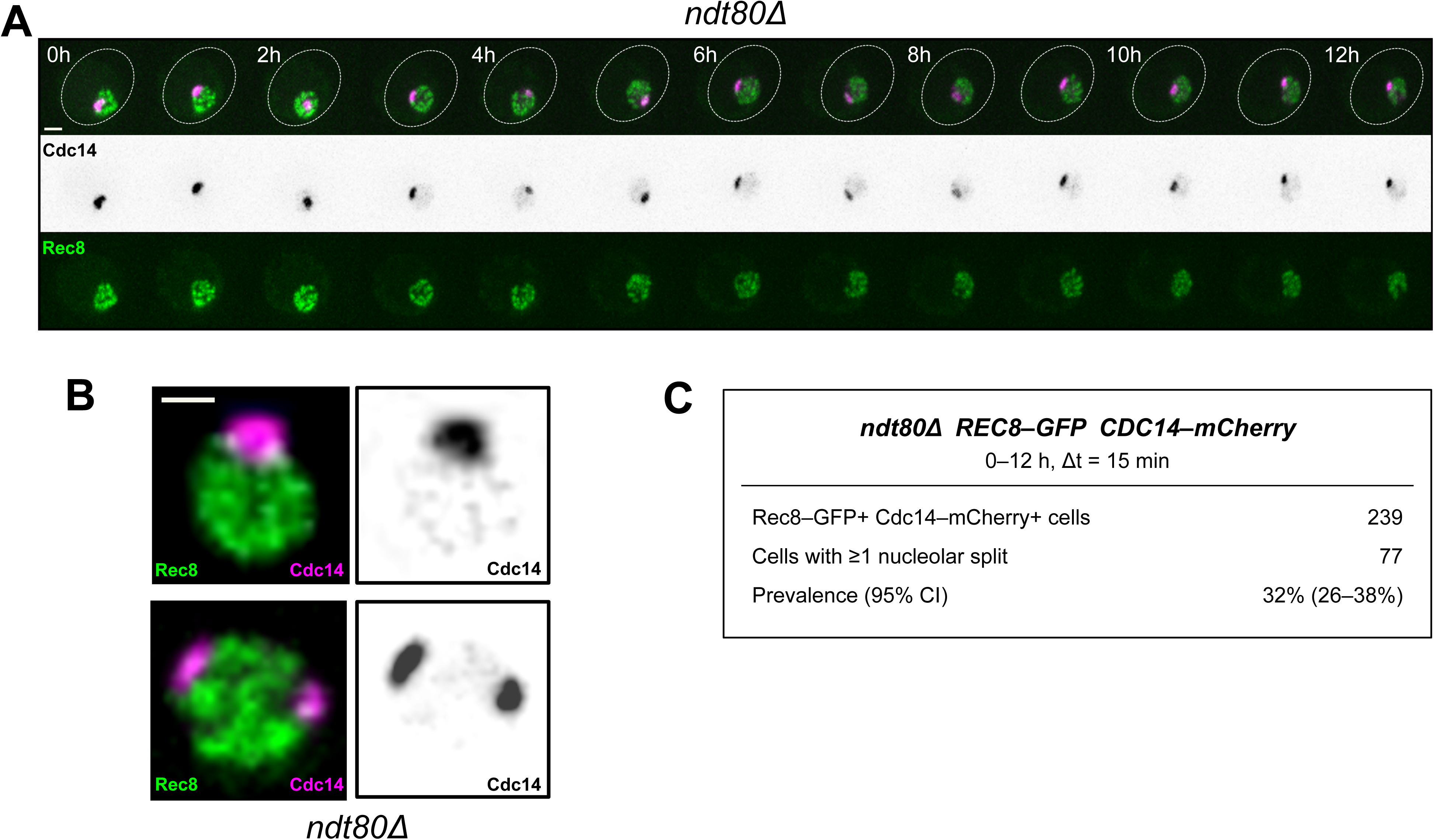
Prophase-I reference in *ndt80Δ*: Cdc14 nucleolar distribution with occasional two-focus episodes (A) Representative time-lapse of an *ndt80Δ* cell (Rec8–GFP gate) showing a predominant single nucleolar Cdc14–mCherry focus with an occasional transition into two spatially separated foci that later re-join within the same series. Micrometric scale bars and frame interval are indicated. (B) Still frames from the same time-lapse highlighting the two-focus episode and re-fusion under the predefined scoring criteria (intensity dip and centroid separation across consecutive 15-min frames). Imaging conditions were matched across figures (Methods). (C) Summary table indicating the number of time-lapse series analysed and the number of *ndt80Δ* cells scored under these criteria, including the subset showing at least one two-focus episode within the nucleolar territory.

### 2.2. Recombination failure increases the two-focus Cdc14 behaviour

Prompted by the initial observation of a two-focus redistribution of Cdc14, we next asked whether this behaviour is influenced by three features of the meiotic recombination landscape: inter-homolog recombination, the accumulation of unrepaired meiotic DSBs, and the outcome of homology search. To isolate a condition in which these three dimensions are simultaneously impacted, we examined *dmc1Δ ndt80Δ*, in which the meiosis-specific recombinase Dmc1 is absent. Time-lapse series in this background showed that the nucleolar Cdc14–mCherry signal frequently resolved into two spatially separated foci within single cells (Fig. 2A–C; Supplementary Movie S2). Quantification restricted to prophase-I cells identified by Rec8–GFP indicated a marked increase in the fraction of cells that displayed the two-focus Cdc14 behaviour relative to the *ndt80Δ* reference (Table 2; see Fig. 2B–C), using the same prophase-I gating and the same event definition as in 2.1 (Methods; Supplementary Table S2), ensuring that differences reflected genotype rather than scoring. Recurrent two-focus episodes were readily observed within single cells in *dmc1Δ ndt80Δ* (Fig. 2A–C).

**Figure 2.**
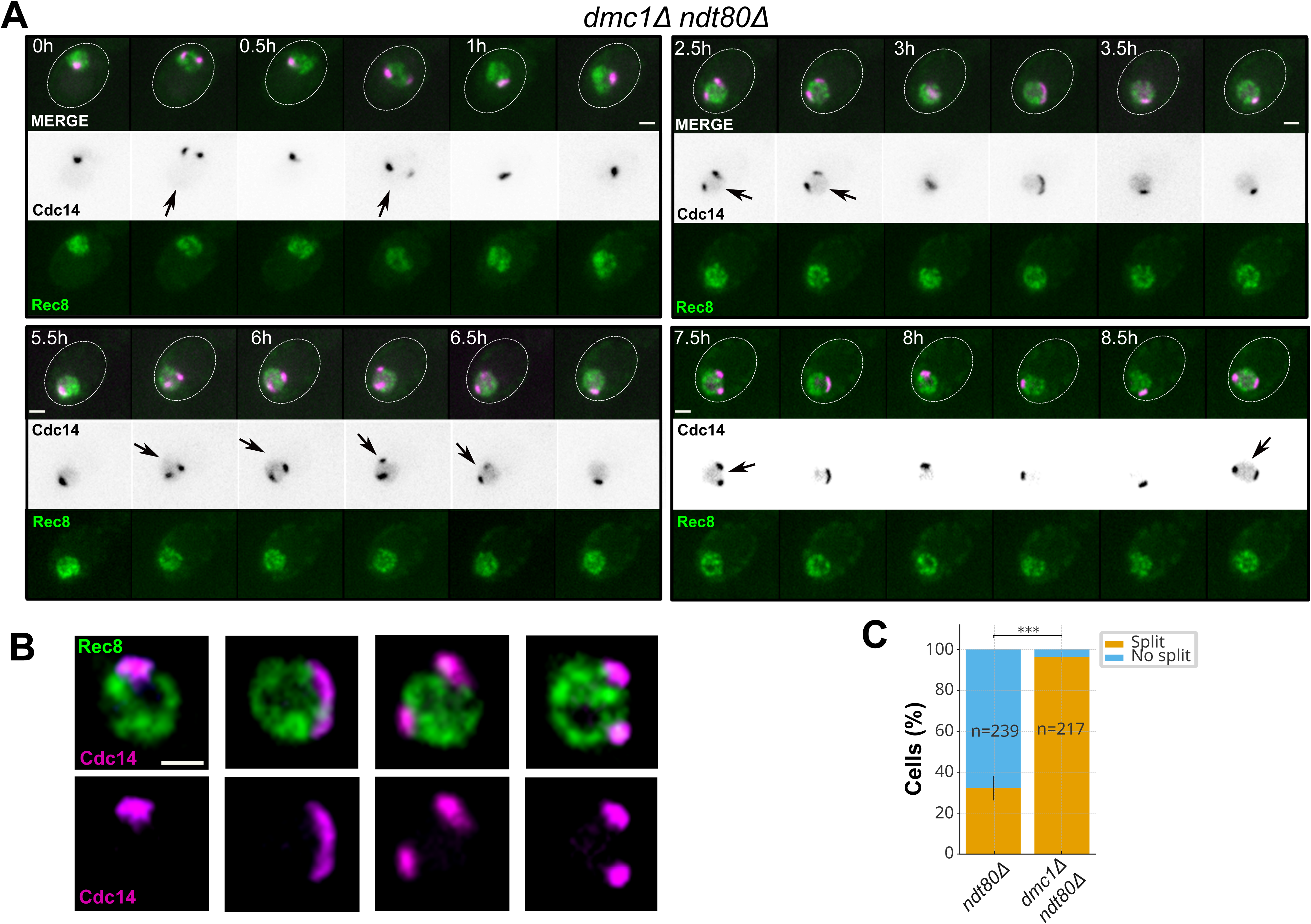
Recombination failure (*dmc1Δ ndt80Δ*) increases the two-focus Cdc14 behaviour (A) Representative time-lapse of a *dmc1Δ ndt80Δ* prophase-I cell (Rec8–GFP gate) showing frequent two-focus episodes of nucleolar Cdc14–mCherry within a single series. Micrometric scale bars and frame interval are indicated. (B) Representative still images of nucleolar Cdc14–mCherry under the same imaging conditions, illustrating cells with a single compact nucleolar focus and cells displaying two spatially separated nucleolar foci. Images are shown at higher magnification than in Fig. 1 to emphasize the intensity dip and spatial separation used in the event definition. Scale bars are indicated. (C) Stacked bar plot showing, for *ndt80Δ* and *dmc1Δ ndt80Δ*, the percentage of Rec8–GFP-positive (prophase-I) cells that did or did not display ≥1 two-focus episode under the predefined scoring criteria (intensity dip plus centroid separation across consecutive 15-min frames, within a ≤12-h window). Corresponding numerical values and statistical tests are summarized in Table 2. Acquisition settings were matched to those used in Fig. 1.

### 2.3. Nucleolar splitting of Cdc14 reveals a dynamic nucleolar territory

To determine whether the two-focus behaviour of Cdc14 reflects a property of the nucleolus rather than a fluorophore-specific artefact or a Cdc14-restricted redistribution, we analysed strains producing Nop56–GFP and Cdc14–mCherry within the same *ndt80Δ*-based time window used for prophase I scoring in the single-nucleolar-label experiments, applying the same predefined event criteria. *NOP56* encodes a core component of box C/D snoRNPs that marks the bulk nucleolus; its multivalent interactions support condensate formation within nucleolar subdomains (liquid–liquid / polymer–polymer phase separation) [23], providing a compartmental reference independent of Cdc14. In this dual-nucleolar labelling setting (Nop56–GFP plus Cdc14–mCherry), two-focus episodes of Cdc14 remained confined to the Nop56-defined nucleolar territory and showed measurable temporal coincidence with Nop56–GFP (Fig. 3A–C; Supplementary Movie S3). Pixel-by-pixel intensity scatter plots confirmed this co-variation: a representative field of view showed a high Pearson correlation (r ≈ 0.85), a compact single nucleolus reached r ≈ 0.90, and the same nucleolus after a splitting episode still displayed a similarly high coefficient (r ≈ 0.87; Fig. 3C). These examples indicate that Cdc14 largely remains within the Nop56-marked compartment even when the nucleolus adopts a two-focus configuration. Because simultaneous imaging of two nucleolar fluorophores increases light dose, a factor known to perturb biomolecular condensates under intensive or prolonged illumination [40, 41], analyses here were limited to compartment identity and temporal coincidence using the same event definition; absolute frequencies were not contrasted to the Rec8–GFP/Cdc14–mCherry datasets used for genotype comparisons, which were acquired with a lower nucleolar light budget. On this basis, we referred to this event as nucleolar splitting. Occasional divergences in timing or amplitude between Cdc14 and Nop56 signals were noted in individual series (Supplementary Movie S3) and are consistent with distinct partitioning across nucleolar subdomains, without altering the compartment-level conclusion [23].

**Figure 3.**
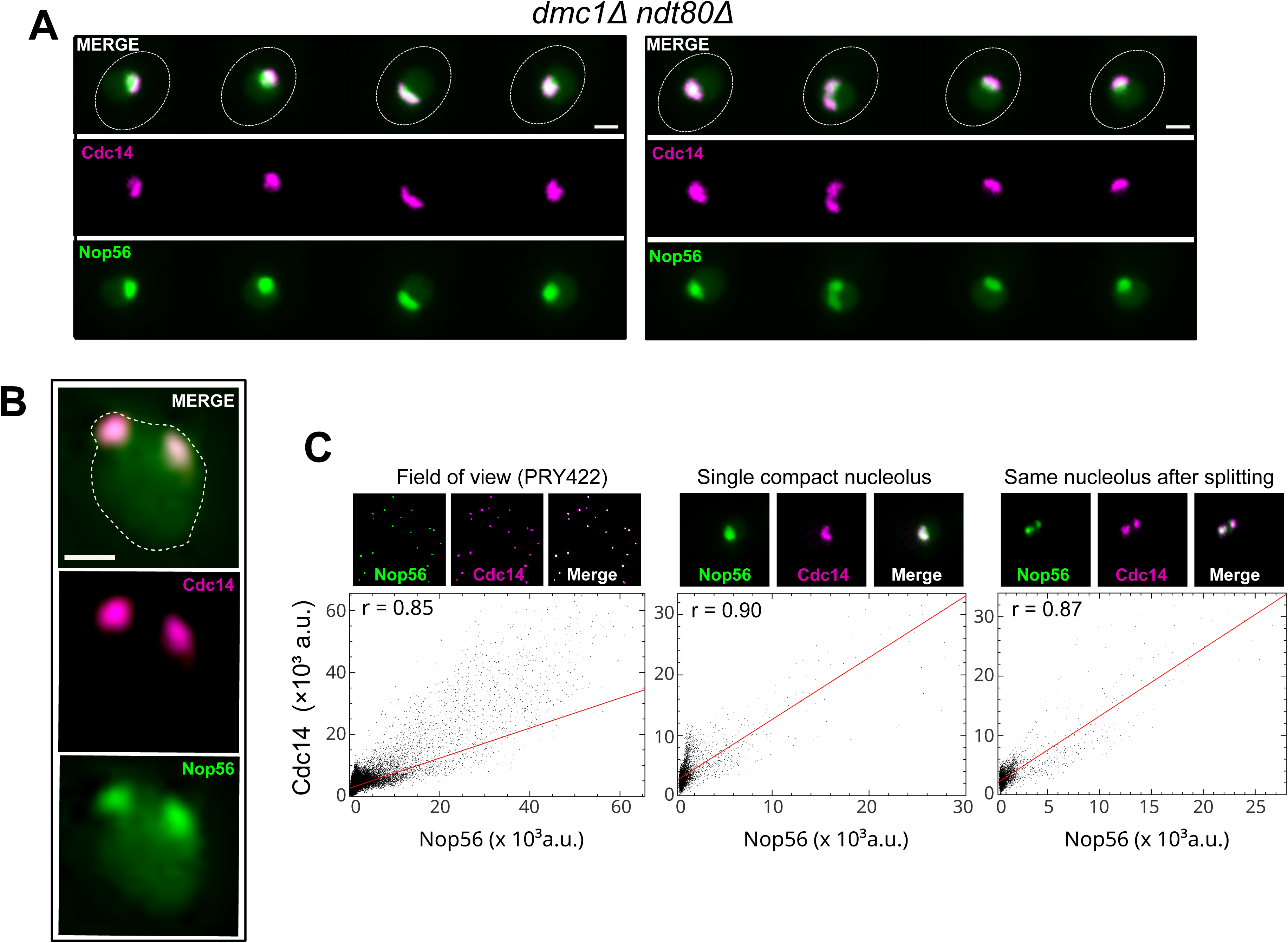
The two-focus Cdc14 behaviour maps to the nucleolus: coincidence with Nop56–GFP (A) Representative time-lapse of a *dmc1Δ ndt80Δ* meiotic cell co-expressing Nop56–GFP and Cdc14–mCherry. The Cdc14–mCherry signal undergoes a transition from a single compact nucleolar focus to two spatially separated foci that remain confined within the Nop56–GFP-defined nucleolar territory before re-fusing. Micrometric scale bars and frame interval are indicated. (B) Single-frame example from a *dmc1Δ ndt80Δ* nucleus co-expressing Nop56–GFP and Cdc14–mCherry. The diffuse Nop56–GFP signal outlines the bulk nucleolus and extends slightly into the nucleoplasm, while Cdc14–mCherry appears in a two-focus configuration within the same nucleolar region, with an intensity valley separating the two peaks. (C) Representative pixel-intensity scatter plots (Cdc14–mCherry versus Nop56–GFP) and the corresponding images used for the analysis, illustrating the degree of spatial co-localisation during two-focus episodes. Pearson correlation coefficients for individual series are indicated for each scatter plot. Because simultaneous imaging of two nucleolar fluorophores increases light dose, this dual-nucleolar configuration was used only to validate compartment identity and co-dynamics, not to compare absolute event frequencies with the datasets in which only Cdc14–mCherry marks the nucleolus (Rec8–GFP + Cdc14–mCherry strains used for genotype comparisons).

### 2.4. Elevated nucleolar splitting persists without meiotic DSB formation

To determine whether the elevated nucleolar splitting observed in *dmc1Δ ndt80Δ* relative to *ndt80Δ* stems from the accumulation of unrepaired DNA double-strand breaks (DSBs) or from deficits in homolog engagement, we abolished meiotic DSB formation using *spo11-y135f*, an allele of *SPO11* that encodes a catalytically inactive Spo11 variant. We introduced *spo11-y135f* into *ndt80Δ* and *dmc1Δ ndt80Δ*, keeping the prophase-I scoring window and the event definition constant, in strains carrying Rec8–GFP as a prophase-I marker together with Cdc14–mCherry to visualize the nucleolar pool. Removing DSB formation eliminates the substrate for resection and presynaptic filament assembly, allowing us to isolate the contribution of Spo11-dependent DSB load. Time-lapse series showed that nucleolar splitting remained frequent in *spo11-y135f dmc1Δ ndt80Δ* and was also elevated in *spo11-y135f ndt80Δ* compared with the *ndt80Δ* reference (Fig. 4A–C; Table 2). Quantification of the prevalence of nucleolar splitting (fraction of Rec8–GFP-positive cells with ≥1 episode) confirmed that both *spo11-y135f* backgrounds showed levels well above *ndt80Δ* (Table 2), whereas the difference between *spo11-y135f ndt80Δ* and *spo11-y135f dmc1Δ ndt80Δ* was modest (Fig. 4C; Table 2). These data indicated that DSB formation per se was not the main driver and that nucleolar splitting of Cdc14 can remain elevated even in the absence of Spo11-dependent DSBs.

**Figure 4.**
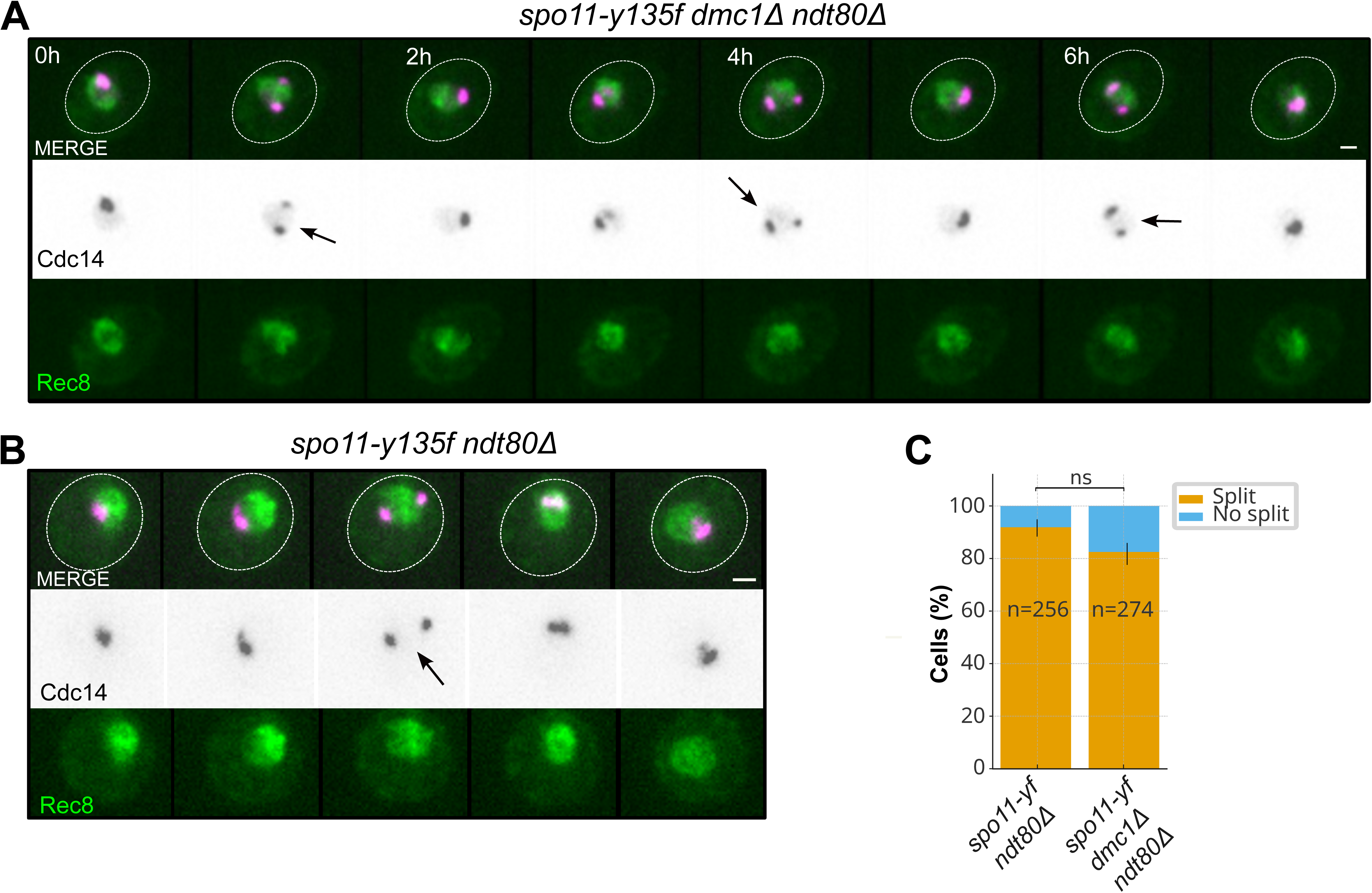
Elevated nucleolar splitting persists without meiotic DSB formation (A) Representative time-lapse of a *spo11-y135f dmc1Δ ndt80Δ* prophase-I cell (Rec8–GFP gate) showing frequent nucleolar splitting episodes of Cdc14–mCherry (two-focus/re-fusion cycles) in the absence of Spo11-dependent DSBs. Micrometric scale bars and frame interval are indicated. (B) Representative time-lapse of a *spo11-y135f ndt80Δ* prophase-I cell (Rec8–GFP gate) imaged under the same conditions, illustrating that nucleolar splitting remains common even when Dmc1 is present and Spo11-dependent DSB formation is abolished. Scale bars as in (A). (C) Stacked bar plot showing, for *spo11-y135f ndt80Δ* and *spo11-y135f dmc1Δ ndt80Δ*, the percentage of Rec8–GFP-positive (prophase-I) cells that did or did not display ≥1 nucleolar splitting episode under the predefined scoring criteria (intensity dip plus centroid separation across consecutive 15-min frames, within a ≤12-h window). Bars or points show counts with 95% confidence intervals as appropriate (Methods). Corresponding numerical values and statistical tests, including comparison to the *ndt80Δ* reference, are summarized in Table 2. Acquisition settings were matched to those used in Figs. 1–3.

### 2.5. Nucleolar splitting is decoupled from the activity state of the meiotic recombination checkpoint

We next asked whether nucleolar splitting parallels the activity state of the meiotic recombination checkpoint at the population level. Population immunoblots for Hop1 phosphorylation and Cdc14–mCherry abundance were acquired on matched 0–10 h time courses in the same *REC8-GFP CDC14-mCherry* backgrounds used for live imaging (*ndt80Δ*, *dmc1Δ ndt80Δ*, *spo11-y135f ndt80Δ* and *spo11-y135f dmc1Δ ndt80Δ*; see Table 1), with Pgk1 as loading control (Fig. 5A–D). Hop1 phosphorylation was weak or transient in *ndt80Δ* and strong and sustained in *dmc1Δ ndt80Δ*, and essentially undetectable in *spo11-y135f* backgrounds, consistent with persistent recombination-checkpoint activation in *dmc1Δ ndt80Δ* and with the abolition of Spo11-dependent DSB formation in *spo11-y135f* strains. Quantification of Hop1 phosphorylation relative to total Hop1 at 6 and 8 h (Fig. 5E) confirmed this pattern, with checkpoint activation high in *dmc1Δ ndt80Δ*, intermediate in *ndt80Δ* and minimal in *spo11-y135f* backgrounds. Cdc14–mCherry levels were broadly comparable across genetic backgrounds and time points (Fig. 5A–D). Yet nucleolar splitting was high in both *dmc1Δ ndt80Δ* and *spo11-y135f* backgrounds and low in *ndt80Δ* (Table 2), indicating that nucleolar splitting did not parallel Hop1 phosphorylation at the population level and appeared decoupled from meiotic recombination-checkpoint activity under the conditions tested. Given the similar Cdc14–mCherry levels, the changes in nucleolar splitting were not explained by alterations in Cdc14 abundance.

**Figure 5.**
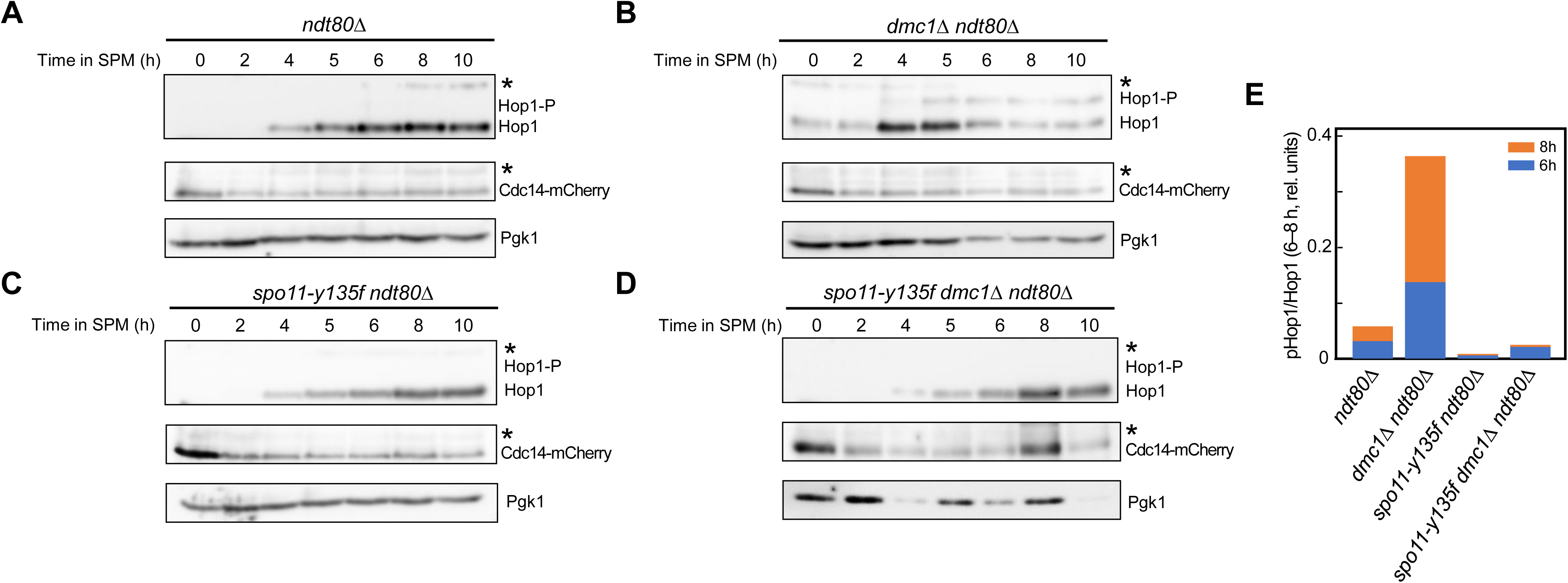
Population checkpoint readouts do not track nucleolar splitting (A) Time-course immunoblot (0–10 h) for *ndt80Δ* showing Hop1, Hop1 phosphorylation (mobility-shifted band), Cdc14–mCherry and Pgk1 as loading control. (B) As in (A), for *dmc1Δ ndt80Δ*, illustrating strong and persistent Hop1 phosphorylation together with stable Cdc14–mCherry levels. (C) As in (A), for *spo11-Y135F ndt80Δ*, illustrating markedly reduced Hop1 phosphorylation across the time course in a background lacking Spo11-dependent meiotic DSB formation. (D) As in (A), for *spo11-Y135F dmc1Δ ndt80Δ*, showing a similarly low Hop1 phosphorylation pattern when both Spo11-dependent DSB formation and Dmc1 are absent. Cdc14–mCherry and Pgk1 signals are shown for all genotypes as controls for Cdc14 abundance and loading. (E) Quantification of Hop1 phosphorylation relative to total Hop1 at 6 and 8 h for the four genotypes. Bars represent average Hop1-P/Hop1 ratios calculated from these time points under the same culture and sampling conditions; corresponding numerical values and details of the quantification procedure are provided in Methods. All time courses were performed in the *REC8–GFP CDC14–mCherry* strains used for live imaging in Figs. 1–4 and 6 (genotypes in Table 1).

**Table 1.** List of yeast strains used in this study.

| Strain | Genotype |
| --- | --- |
| PRY414 | *MAT a/α ho::LYS2/hisG, ndt80::hphMX6/", CDC14-mCherry-ClonNat/", REC8-GFP::LEU2::KanMX4/", trp1/TRP1, his4X::LEU2 (NgoMIV; ori)-URA3/HIS4* |
| PRY221 | *MAT a/α ho::LYS2/ho::hisG, ndt80::hphMX6/", CDC14-mCherry-ClonNat/", REC8-GFP::LEU2::KanMX4/", his4/", dmc1Δ::KanMX4/", ura3/"* |
| PRY416 | *MAT a/α ho::hisG/", ndt80::hphMX6/", CDC14-mCherry-ClonNat/", REC8-GFP::LEU2::KanMX4/", trp1/", his4X::LEU2 (NgoMIV; ori)-URA3/", arg4/", spo11-Y135F-HA-URA3/", dmc1Δ::KanMX4/DMC1* |
| PRY346 | *MAT a/α, ho::LYS2, lys2, ndt80::hphMX6/", CDC14-mCherry-ClonNat/", REC8-GFP::LEU2::KanMX4/", dmc1Δ::KanMX4/", spo11-Y135F-HA-URA3/", arg4/", HIS4/his4* |
| PRY420 | *MAT a/α NOP56-GFP::URA3, CDC14-mCherry::ClonNat, ndt80::hphMX6, leu2/", lys2/", his4/", trp1/"* |
| PRY422 | *MAT a/α NOP56-GFP::URA3, CDC14-mCherry::ClonNat, ndt80::hphMX6, dmc1Δ::KanMX4, leu2/", lys2/", his4/", trp1/"* |

### 2.6. Timing and persistence of nucleolar splitting across recombination contexts

To assess persistence rather than onset, we used three complementary timing metrics: (i) the distribution of all individual event times (Fig. 6A), (ii) the fraction of cells with ≥1 event beyond predefined thresholds (6.0 h and 9.0 h; Fig. 6B; Table S2), and (iii) for each cell, the time of its last observed event (TLAST; Table S3). The 6.0 h and 9.0 h thresholds were selected to balance late-time information with limits on cumulative illumination.

**Figure 6.**
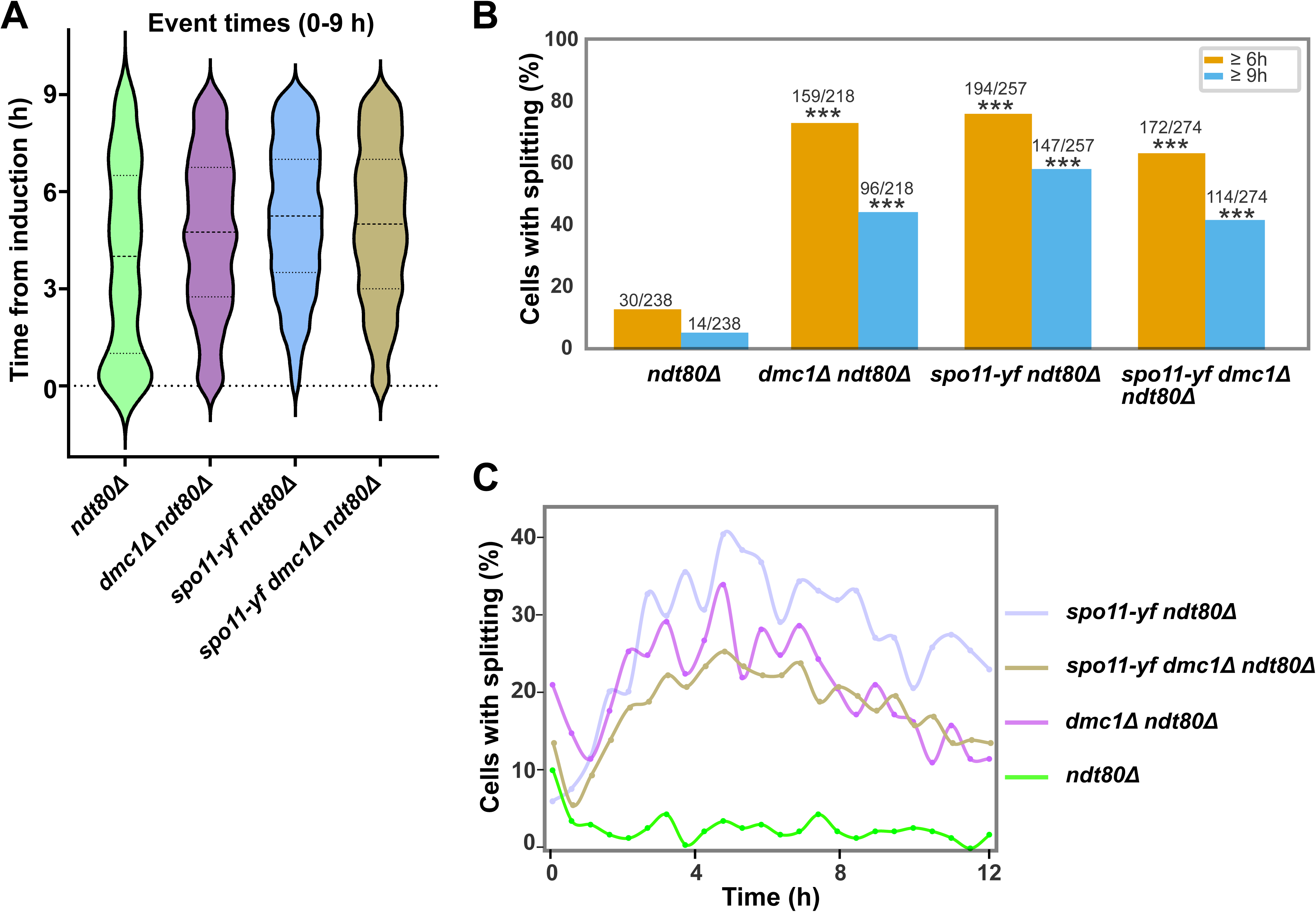
Temporal persistence of splitting across genetic backgrounds (A) Violin plots showing the distribution of all nucleolar splitting event times (frames with ≥1 splitting event) from 0 to 9.0 h after induction for *ndt80Δ*, *dmc1Δ ndt80Δ*, *spo11-y135f ndt80Δ* and *spo11-y135f dmc1Δ ndt80Δ*. Central lines indicate median event times, with the width of each violin reflecting the density of events. Event-time quartiles (Q1/median/Q3, in hours) were 1.0/4.0/6.5 for *ndt80Δ*, 2.75/4.75/6.75 for *dmc1Δ ndt80Δ*, 3.5/5.25/7.0 for *spo11-Y135F ndt80Δ* and 3.0/5.0/7.0 for *spo11-y135f dmc1Δ ndt80Δ*. The number of events contributing to each violin was 157, 1075, 1598 and 1173, respectively (Table S3). (B) Fraction of cells with ≥1 nucleolar splitting event at or after 6.0 h and at or after 9.0 h for the same genotypes. Bars report percentages, and numbers above each bar indicate n/N cells with at least one late event under the corresponding threshold. Asterisks on mutant bars denote significant increases relative to *ndt80Δ* at the same threshold (2×2 χ² tests, df = 1; χ² = 86.7–197.2, p < 10⁻²⁰; full statistics in Table S2). (C) Temporal profiles of nucleolar splitting activity. Lines show the fraction of cells with ≥1 event per 0.5 h time bin over the 12 h time-course. Activity peaks early and remains low in *ndt80Δ*, whereas *dmc1Δ* and *spo11-Y135F* backgrounds display broader, sustained activity extending into mid–late time bins. A χ² test over the full binned time-course confirms a significant difference between *ndt80Δ* and *dmc1Δ ndt80Δ*, while *spo11-Y135F ndt80Δ* and *spo11-Y135F dmc1Δ ndt80Δ* are statistically indistinguishable (Table S2). TLAST summaries and Kruskal–Wallis tests for last-event times per cell are provided in Table S3.

The violin distributions of all events scored up to 9.0 h (Fig. 6A) and the per-cell persistence table (Table S2) showed a graded pattern: in *ndt80Δ* the phenomenon concentrates early and wanes, whereas it remains frequent at late times in *dmc1Δ* and *spo11* contexts. Quantitatively, at 6.0 h (25 frames) late-event fractions were ∼12.6% in *ndt80Δ*, ∼72.9% in *dmc1Δ ndt80Δ*, ∼75.5% in *spo11-y135f ndt80Δ*, and ∼62.8% in *spo11-y135f dmc1Δ ndt80Δ*, supporting marked heterogeneity across genotypes; the same ordering was observed at the very-late 9.0 h cut-off (Fig. 6B; Table S2). Two-by-two χ² tests comparing each mutant to *ndt80Δ* at the same threshold confirmed that all contrasts were highly significant (χ² = 86.7–197.2, df = 1, p < 10⁻²⁰; full statistics in Table S2). Consistently, per-cell last-event medians (TLAST; Table S3) were earliest in *ndt80Δ* and shifted later in *dmc1Δ* and *spo11* backgrounds, in agreement with Kruskal–Wallis tests on TLAST distributions (e.g., *ndt80Δ* vs *dmc1Δ ndt80Δ*, χ² = 72.629, df = 1, p < 2.2×10⁻¹⁶; Table S3). To avoid reliance on a single cut-off and to visualize when the population is active, a bin-based analysis (0.5 h bins; Fig. 6C) showed that the proportion of cells with ≥1 event per bin peaks early and remained low in *ndt80Δ*, whereas *dmc1Δ* and *spo11* backgrounds displayed broader activity extending into mid–late time bins; a χ² test over the full time-course confirmed a significant difference between *ndt80Δ* and *dmc1Δ ndt80Δ*, while *spo11-Y135F ndt80Δ* and *spo11-Y135F dmc1Δ ndt80Δ* were statistically indistinguishable (Table S2). Collectively, these timing analyses indicated that splitting persisted into late prophase in recombination-defective and DSB-free settings, but declined in the *ndt80Δ* reference under identical scoring conditions.

## 3. Discussion

We identify a transient two-focus redistribution of Cdc14 within the nucleolus during meiotic prophase I and show that its prevalence varies with the recombination landscape under a common prophase-anchored scoring regime. The behaviour is infrequent in a recombination-competent reference (*ndt80Δ*), increases when recombination fails (*dmc1Δ*), and remains elevated when Spo11-dependent DSB formation is abolished (*spo11-y135f*). Validation with Nop56-GFP places the event within the nucleolar compartment, and genotype-matched acquisition/analysis rules support that these differences reflect biology rather than imaging bias.

We use the term “nucleolar splitting” strictly for two-focus transitions that satisfy the predefined intensity-dip and separation criteria (Methods), and only after compartment validation with Nop56-GFP. The term does not imply rDNA breakage or irreversible nucleolar disruption; it denotes a reversible partition of the nucleolar signal, consistent with condensate-like behaviour [23, 42].

Against this background, we next asked whether bulk recombination-checkpoint readouts track the phenomenon at the population level. Population checkpoint readouts do not map onto the pattern of nucleolar splitting. Strong Hop1 phosphorylation in *dmc1Δ*, weak or transient phosphorylation in *ndt80Δ*, and undetectable phosphorylation in *spo11–y135f* coexist with high splitting in *dmc1Δ* and *spo11–y135f* and low splitting in *ndt80Δ* (Figs. 5A–B; Table 2; see also Figs. 2, 4, and 6A). Under the present scoring conditions, the phenomenon appears decoupled from recombination-checkpoint activity at the population level [10, 43, 44].

**Table 2.** Prevalence of nucleolar splitting under Rec8–GFP prophase-I gating.

| Genotype | Total cells (N) | Cells with $\geq 1$ splitting event (n) | Prevalence (%) |
| --- | --- | --- | --- |
| *ndt80Δ* | 238 | 74 | 31.1 |
| *dmc1Δ ndt80Δ* | 218 | 209 | 95.9 |
| *spo11-Y135F ndt80Δ* | 257 | 236 | 91.8 |
| *dmc1Δ spo11-y135f ndt80Δ* | 274 | 225 | 82.1 |

Event prevalence was defined as the fraction of cells with $\geq 1$ nucleolar splitting episode over the 0–12 h time-course under Rec8–GFP prophase-I gating (Methods 4.4).

Although bulk checkpoint markers do not track the behaviour, an upstream ATR/ATM input may still contribute in specific contexts. During persistent Mec1 activation, as in the *dmc1Δ* background, Mec1-dependent phosphorylation of Zip1 at centromeres dismantles Zip1-mediated centromere coupling [12, 45]. Loss of coupling has been associated with altered whole-chromosome dynamics, which in our framework could facilitate force transmission to the rDNA and thereby favour nucleolar splitting. We therefore view this pathway as one plausible contributing route rather than a sole driver.

In *spo11–y135f*, a complementary DSB-independent route is plausible. Zip1-mediated centromere coupling and pairing can persist and channel force transmission in the absence of breaks, while telomere bouquet formation and telomere-led rapid prophase movements (RPM) proceed through the Ndj1–Mps3–Csm4 LINC pathway: bouquet sets the attachment geometry, RPM supplies the driving forces. Together, these features increase the degrees of freedom of whole-chromosome motion and facilitate force transfer to the rDNA [15, 18, 19]. In future work, genetic perturbations that differentially affect telomere attachment versus force production could be used to challenge this model.

We interpret the two-focus behaviour as more than a passive consequence of bulk chromosomal motion. The Nop56–GFP dataset shows that the event involves the nucleolus as a compartment, while occasional divergences from Cdc14–mCherry indicate that nucleolar subdomains need not move in perfect register. Work in mitotic cells already supports a mixed-condensate architecture at the rDNA locus. Quantitative imaging showed that Cdc14 and other rDNA-bound factors occupy a compressible polymer condensate, whereas Nop56-marked ribonucleoproteins form a more homogeneous, liquid-like phase that neither compacts when rDNA loops contract nor disperses when the rDNA array is lost [23]. In physical terms, the nucleolus behaves as a biomolecular condensate [22, 46]; transient splitting then reflects a rheological response of that condensate to chromosome-borne forces when homolog engagement is compromised or delayed [15, 47, 48]. The rapid return to a single body is naturally explained by coalescence/coarsening dynamics once the deforming constraint relaxes [6].

This framework is consistent with our genetics. Elevated splitting in *dmc1Δ* and in *spo11–y135f*—despite opposite DSB contexts—implicates the state of homolog engagement rather than DSB load as the decisive variable [49]. In budding yeast, the rDNA array on chromosome XII does not assemble synaptonemal complex [18, 25–28]; accordingly, forces arising from pairing attempts and centromere/telomere reorganizations during prophase I can act without SC reinforcement at the rDNA, making the nucleolus a sensitive readout of that mechanical landscape [15, 18, 47]. The lack of correspondence with checkpoint readouts further supports a mechanical, rather than signalling-driven, origin under our conditions [10].

Finally, the modest desynchrony sometimes observed between Cdc14–mCherry and Nop56–GFP fits a model in which Cdc14-rich and Nop56-rich subcompartments differ in composition and relaxation times (liquid-like versus more polymer-dominated features), allowing small phase-specific offsets in their motions without altering the compartment-level conclusion [23]. In sum, “nucleolar splitting” provides a mesoscale, mechanical readout of prophase-I chromosome dynamics, and its fast resolution reflects the material properties of the nucleolus itself.

DSB-independent routes of homolog proximity likely contribute to this picture. Prior work described non-homologous centromere coupling in early prophase I and its dependence on the synaptonemal complex component Zip1, which can organize centromeres in the absence of Spo11-generated breaks [12, 18, 45]. Such coupling may provide ordered coalescence of centromere-associated factors and, in pathological regimes, promote aggregate-like assemblies [50–52]. Within this framework, Zip1-mediated centromere coupling in *spo11–y135f* could sustain pairing attempts and associated force transmission even without DSBs, thereby favouring transient partitioning of the nucleolar condensate; once constraints relax, coarsening would rapidly restore a single body. This hypothesis is consistent with the elevated splitting observed in *spo11–y135f* and can be further refined by future genetic dissection of centromere-coupling pathways [12].

These elements are summarized in a working model (Fig. 7) that links recombination status, homolog engagement, and telomere-driven forces to nucleolar behaviour. In a DSB-free configuration (very early *ndt80Δ* or *spo11-y135F*), telomeres anchored via Ndj1–LINC and Zip1-mediated centromere coupling act on poorly engaged chromosome XII arms, favouring repeated splitting of the Cdc14/Nop56 condensate. When Spo11-dependent DSBs form but interhomolog repair is compromised (for example in *dmc1Δ ndt80Δ*), Mec1-dependent regulation of centromeric Zip1 releases non-homologous couplings but homolog engagement remains defective, so telomere-led forces still fragment the nucleolus. Once Dmc1-dependent interhomolog repair and ZMM/SC assembly establish robust homolog engagement (*NDT80*-proficient or late *ndt80Δ* cells), telomere attachments persist but forces are buffered by paired bivalents and nucleolar splitting subsides.

**Figure 7.**
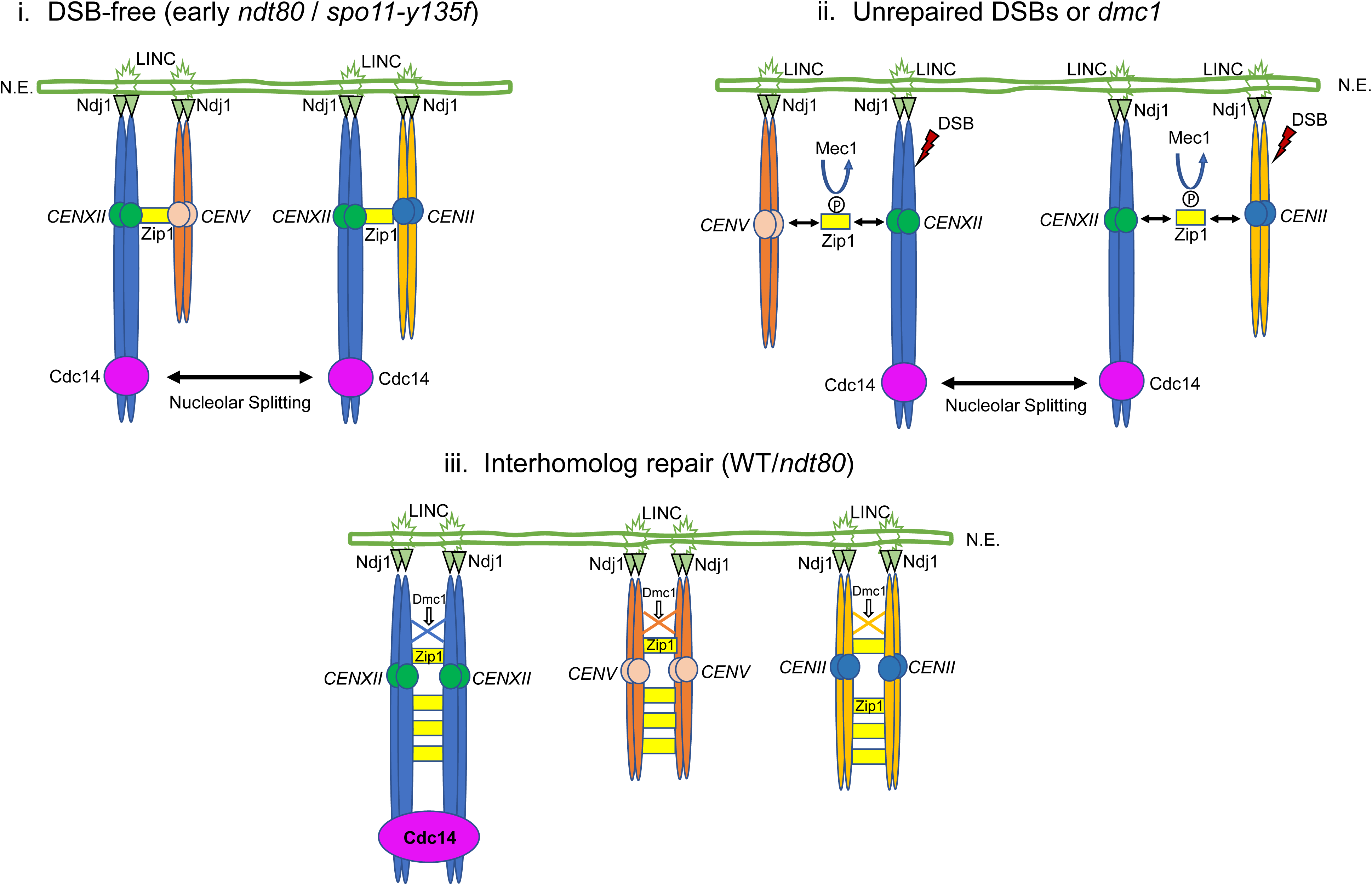
Working model linking recombination status, telomere driven forces, and nucleolar splitting (i) DSB free (early *ndt80Δ* / *spo11-y135F*). Telomeres of homologous chromosomes are anchored to the nuclear envelope (N.E.) through Ndj1–LINC complexes. Before Spo11 dependent DSB formation, Zip1 accumulates at centromeres and promotes non homologous centromere coupling (examples shown for *CENXII* and *CENV*, and for *CENXII* and *CENII*). Under this configuration, rapid prophase movements transmitted from Ndj1–LINC to poorly engaged homologs deform the rDNA bearing region and promote dynamic nucleolar splitting, visualized as oscillation of Cdc14 between two separated nucleolar bodies. (ii) Unrepaired DSBs or *dmc1Δ*. Spo11 induced DSBs activate Mec1, which phosphorylates centromeric Zip1 and releases non homologous centromere couplings. In *dmc1Δ*, or in early *ndt80Δ* before interhomolog repair completes, homolog engagement remains defective despite centromere decoupling. Telomere driven forces persist in this configuration and continue to pull on incompletely paired chromosome XII arms, so nucleolar splitting remains frequent and long lived. (iii) Interhomolog repair (WT / *ndt80*). When Dmc1 driven interhomolog repair and synaptonemal complex assembly are established, homologous centromeres and arms are fully engaged by Zip1. Ndj1–LINC still anchors telomeres, but forces are now buffered by a rigid bivalent, so the nucleolar condensate stabilizes and Cdc14 concentrates in a single nucleolar body with rare splitting episodes. In all panels, only one telomere attachment per homolog and a subset of centromeres are shown for simplicity; Ndj1–LINC anchoring is assumed to operate in all genotypes.

Together, the data and framework converge on a simple working view: under matched scoring conditions, nucleolar splitting reports the state of homolog engagement more than the absolute burden of Spo11-dependent DSBs. The checkpoint does not map onto the behaviour at the population level under our conditions, whereas routes that alter attachment/force routing (centromere coupling via Zip1; telomere-led RPM via the Ndj1–Mps3–Csm4 LINC axis) offer specific, testable predictions for late-time persistence profiles. A next step will be to define how Cdc14 responds to these mechanical inputs in real time, and whether its nucleolar redistribution has consequences for recombination factors at chromosome arms, centromeres, or the rDNA-proximal region. Such experiments may reveal additional modes of Cdc14 control over meiotic recombination that are not apparent from bulk checkpoint readouts alone.

## 4. Materials and Methods

### 4.1. Yeast strains and media

All experiments used Saccharomyces cerevisiae. Datasets underlying Results 2.1, 2.2, 2.4, 2.5 and 2.6 carried Rec8–GFP as a nuclear/chromosome marker together with Cdc14–mCherry to visualize the nucleolar Cdc14 pool. The dual-nucleolar validation in Result 2.3 employed Nop56–GFP plus Cdc14–mCherry and did not include Rec8–GFP. Full genotypes and sources are provided in Table 1. Standard YPD, pre-sporulation, and sporulation media were used as summarized in Supplementary Table S4.

### 4.2. Meiosis induction and time courses

Cells were grown in pre-sporulation medium, shifted to sporulation medium, and imaged at 30 °C under sealed chambers. Time courses started upon transfer to sporulation medium and typically extended up to ∼12 h with a sampling interval of 15 min per frame. Analyses were restricted to prophase-I cells gated by Rec8–GFP where applicable. Acquisition windows and thresholds used to limit cumulative light exposure are specified below and in Result 2.6.

### 4.3. Live-cell microscopy and acquisition parameters

Time-lapse imaging was performed on a spinning-disk confocal platform Roper Scientific with Olympus IX81 inverted microscope; the optical components employed was the 100X 1-4 NA objective and an Evolve (Photometrix) camera. For each time point, a short Z-stack was acquired (typically 11 planes with a Z-step of 0.4 µm, spanning ∼4 µm in depth) around the brightfield focal plane, which approximates the mid-plane of most cells in the field. Under these conditions, the nuclear volume was generally encompassed within the imaged Z-range. Z-stacks were rendered by maximum-intensity projection with fixed LUTs and exposure ranges within each genotype-matched experiment. Excitation, exposure, EM gain/laser power, Z-step and frame size were kept constant per dataset; numerical ranges are reported in Table 4 and Table S1. Time-lapse series in which the Cdc14–mCherry or Nop56–GFP nucleolar signal approached or touched the top or bottom limits of the stack, or appeared clearly truncated in Z, were excluded from analysis.

**Table 3.**
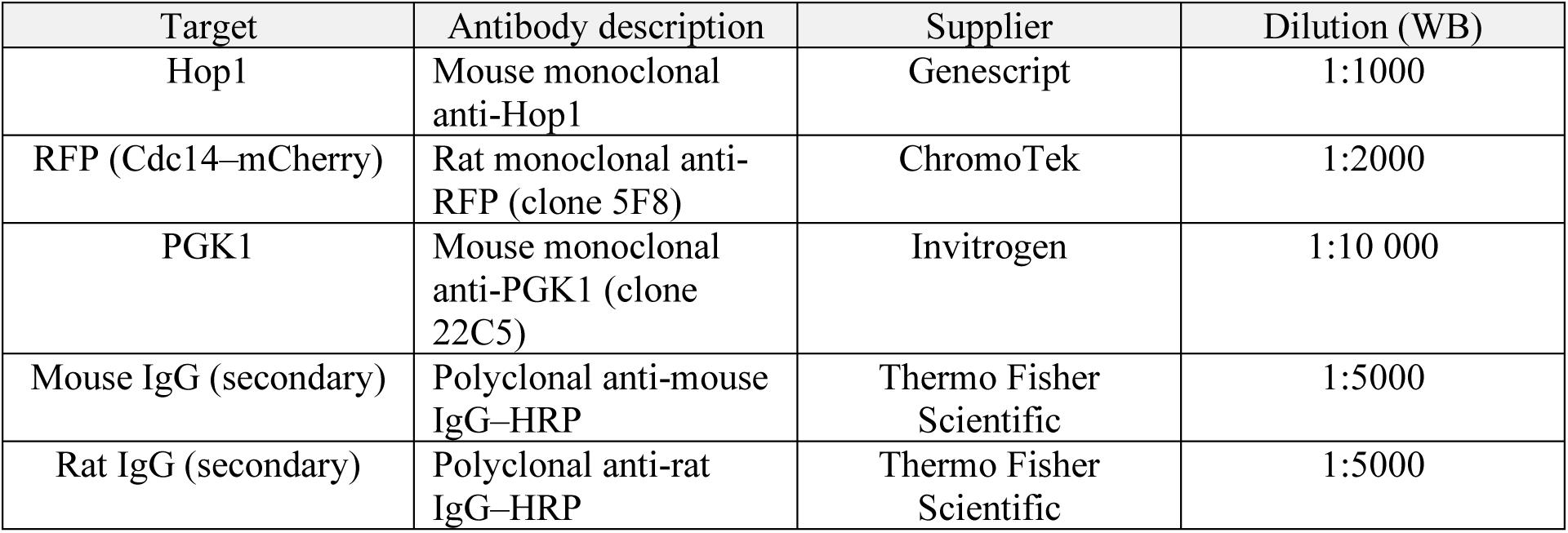
Antibodies used in this study.

| Target | Antibody description | Supplier | Dilution (WB) |
| --- | --- | --- | --- |
| Hop1 | Mouse monoclonal anti-Hop1 | Genescript | 1:1000 |
| RFP (Cdc14–mCherry) | Rat monoclonal anti-RFP (clone 5F8) | ChromoTek | 1:2000 |
| PGK1 | Mouse monoclonal anti-PGK1 (clone 22C5) | Invitrogen | 1:10 000 |
| Mouse IgG (secondary) | Polyclonal anti-mouse IgG–HRP | Thermo Fisher Scientific | 1:5000 |
| Rat IgG (secondary) | Polyclonal anti-rat IgG–HRP | Thermo Fisher Scientific | 1:5000 |

**Table 4.** Channel-specific imaging settings.

| Channel | Z-stack | Exposure time (ms) | Camera readout (MHz) | Laser power (% of max) | Analog gain (×) | EM gain |
| --- | --- | --- | --- | --- | --- | --- |
| TRANS | No | – | 5 | – | 3 (4×) | 800 |
| GFP | Yes (11 × 0.4μm) | 100 | 5 | 18 | 3 (4×) | 800 |
| mCherry | Yes (11 × 0.4μm) | 200 | 5 | 8 | 3 (4×) | 800 |

Live-cell microscopy settings. Channel-specific exposure and laser settings on the EM-CCD

### 4.4. Image processing, gating and event definition

Raw image stacks were processed in Fiji/MetaMorph using a fixed pipeline: drift correction when needed, projection, background estimation from local annuli, and nucleolar ROI tracking from the Cdc14–mCherry signal. Prophase-I gating relied on Rec8–GFP in single-nucleolar datasets. A “two-focus event” was scored in any 15-min frame in which the Cdc14–mCherry signal within the nucleolar ROI displayed two intensity maxima separated by a clear local minimum and a measurable distance between their centroids, with both foci fully contained within the imaged Z-stack; frames were scored only within the first 12 h of the time course to limit cumulative illumination. Exact thresholds, ROI geometry and exclusion rules are summarized in Supplementary Fig. S1A. The term “nucleolar splitting” is used only after compartment validation with Nop56–GFP (Result 2.3) for events that meet these criteria. In dual-nucleolar datasets (Nop56–GFP + Cdc14–mCherry), coincidence was quantified within the NOP56-defined nucleolar territory using time-aligned overlays and line-profile analyses [53]; absolute frequencie were not contrasted to single-label datasets due to different illumination budgets. Visual inspection of representative raw stacks confirmed that these two-focus episodes reflect a physical partition of the nucleolar signal rather than incomplete sampling of elongated nucleolar profiles at the periphery of the Z-stack.

### 4.5. Immunoblotting

Protein extracts were prepared by TCA precipitation [54], resolved by SDS–PAGE [55], transferred to PVDF and probed with antibodies listed in Table 3. Time courses covered 0–10 h under the same sporulation regime used for imaging. Detection used standard ECL within the linear range; exposures were matched across genotypes within an experiment.

### 4.6. Quantification and statistics

Event prevalence was defined as the fraction of cells with ≥ 1 event per genotype; recurrence was the number of events per cell. Timing distributions combined all event times into violin plots and, per cell, the last observed event time (TLAST). Late-event fractions were computed beyond predefined thresholds (primary 6.0 h and 9h) with identical frame sampling across genotypes. Proportions were compared by χ² tests; non-parametric tests were used for timing distributions. Two-sided α = 0.05 was applied, and 95% confidence intervals were obtained by bootstrap when appropriate. Exact n and test values appear in figure panels or Supplementary Tables S2–S3. Analyses were carried out in R and Fiji/MetaMorph with scripts/macros available upon request.

### 4.7. Reporting and data availability

Figure panels show representative series acquired under genotype-matched settings; scale bars and time stamps are indicated where relevant. Source data (per-cell counts, event times, TLAST, and binned activity curves) together with analysis scripts will be provided as Supplementary Fig. S1 and Table S2 and S3.

## Acknowledgements

We thank our colleagues at the Institute of Functional Biology and Genomics (IBFG) for helpful discussions and technical assistance, and the IBFG Microscopy Facility for expert support with live-cell imaging. We are especially grateful to Jesús Pinto-Cruz for assistance with image analysis and statistical treatment of the data. This work was supported partly by the University of Salamanca Master’s Programme in Cellular and Molecular Biology and by MICIU/AEI–ERDF/EU grant PID2021-125830NB-I00.

## Supplementary Figure legends

Supplementary Figure S1. Workflow for nucleolar splitting quantification and persistence metrics.

(A) From time-lapse series, Rec8–GFP-positive prophase-I cells are selected and the nucleolar region is tracked based on Cdc14–mCherry. Two-focus episodes are scored when the nucleolar Cdc14 signal resolves into two intensity maxima separated by a local minimum that satisfies predefined persistence and separation criteria (Methods). (B) For each cell, the times of all scored events are compiled and the last observed event time (TLAST) is extracted. (C) Late-event fractions are obtained as the proportion of cells with ≥ 1 event at or after 6.0 h and at or after 9.0 h under identical frame sampling across genotypes. (D) Population activity curves are generated by grouping events into 0.5 h bins and computing, for each bin, either the fraction of cells with ≥ 1 event (main-text Fig. 6C) or the fraction of total events per genotype (Supplementary Fig. S2). Together, these metrics provide complementary views of nucleolar splitting prevalence and temporal persistence under matched scoring conditions.

Supplementary Figure S2. Alternative summary views of nucleolar-splitting timing profiles.

(A) Cumulative incidence of cells with ≥1 event over the 0–12 h time course for each genotype (fraction of cells that have experienced at least one event by time t). (B) Per-bin distribution of events in 0.5 h intervals, expressed as the percentage of each genotype total, highlighting how events are temporally redistributed rather than simply delayed. (C) Conditional persistence curves (“survival” plots) showing, among event-positive cells, the fraction that still display ≥1 event after time t. (D) Late-persistence progression: for each genotype and threshold time, fraction of cells with ≥1 event at or beyond that threshold. Together, these complementary representations illustrate that nucleolar splitting concentrates early and wanes in *ndt80Δ*, while it remains broadly distributed into mid–late time windows in *dmc1Δ* and *spo11-Y135F* backgrounds.

Supplementary Videos. Additional dual-nucleolar examples for Nop56–GFP/Cdc14–mCherry.

Movies: *ndt80Δ* and *dmc1Δ* representative movies illustrating compartment confinement and occasional timing offsets.

