## Supplemental Figures and tables for "Nucleolar Cdc14 Splitting Reflects Recombination Context and Meiotic Chromosome Dynamics"

#### Supplementary Figure S1

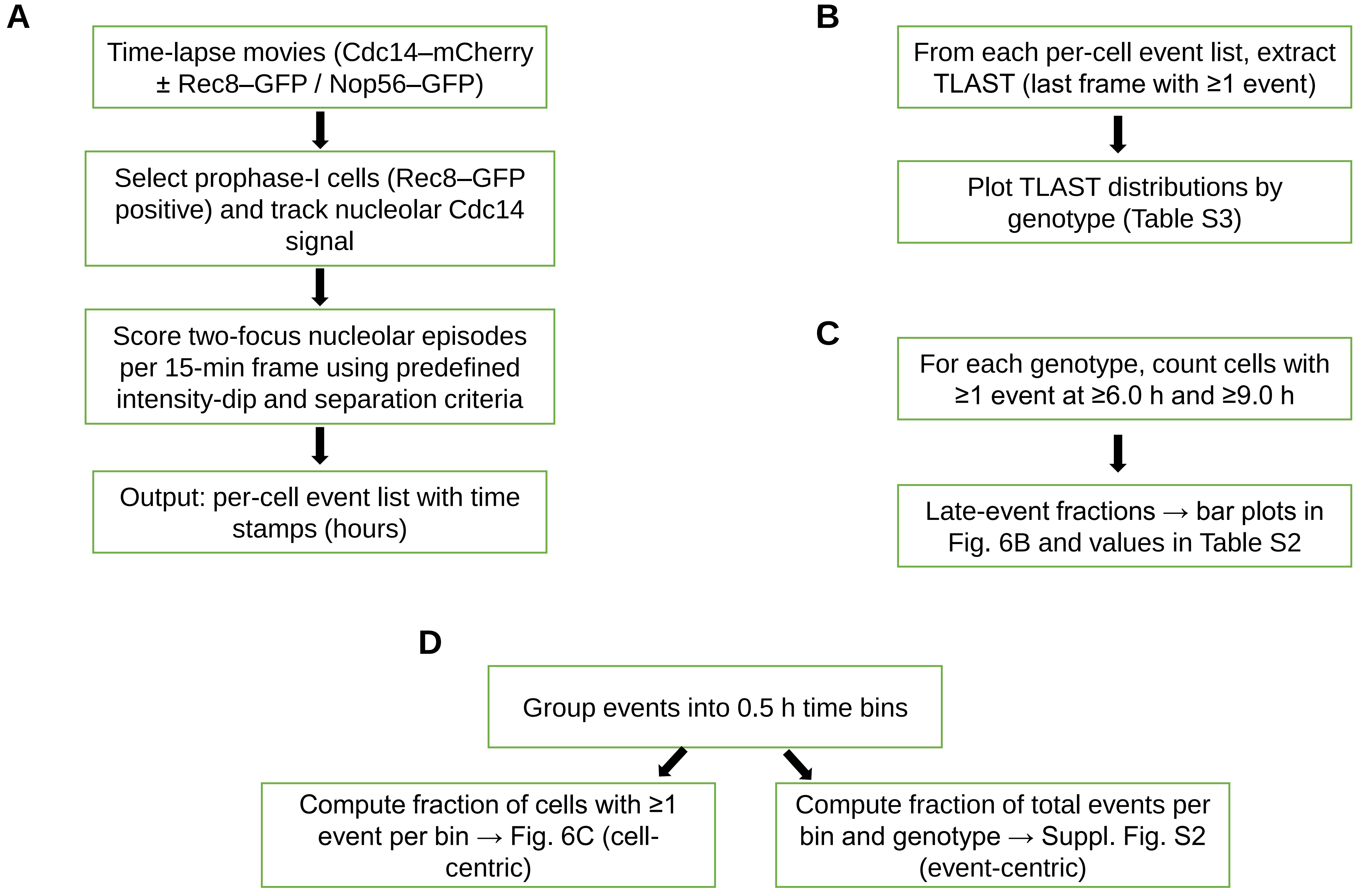

### Supplementary Figure S2

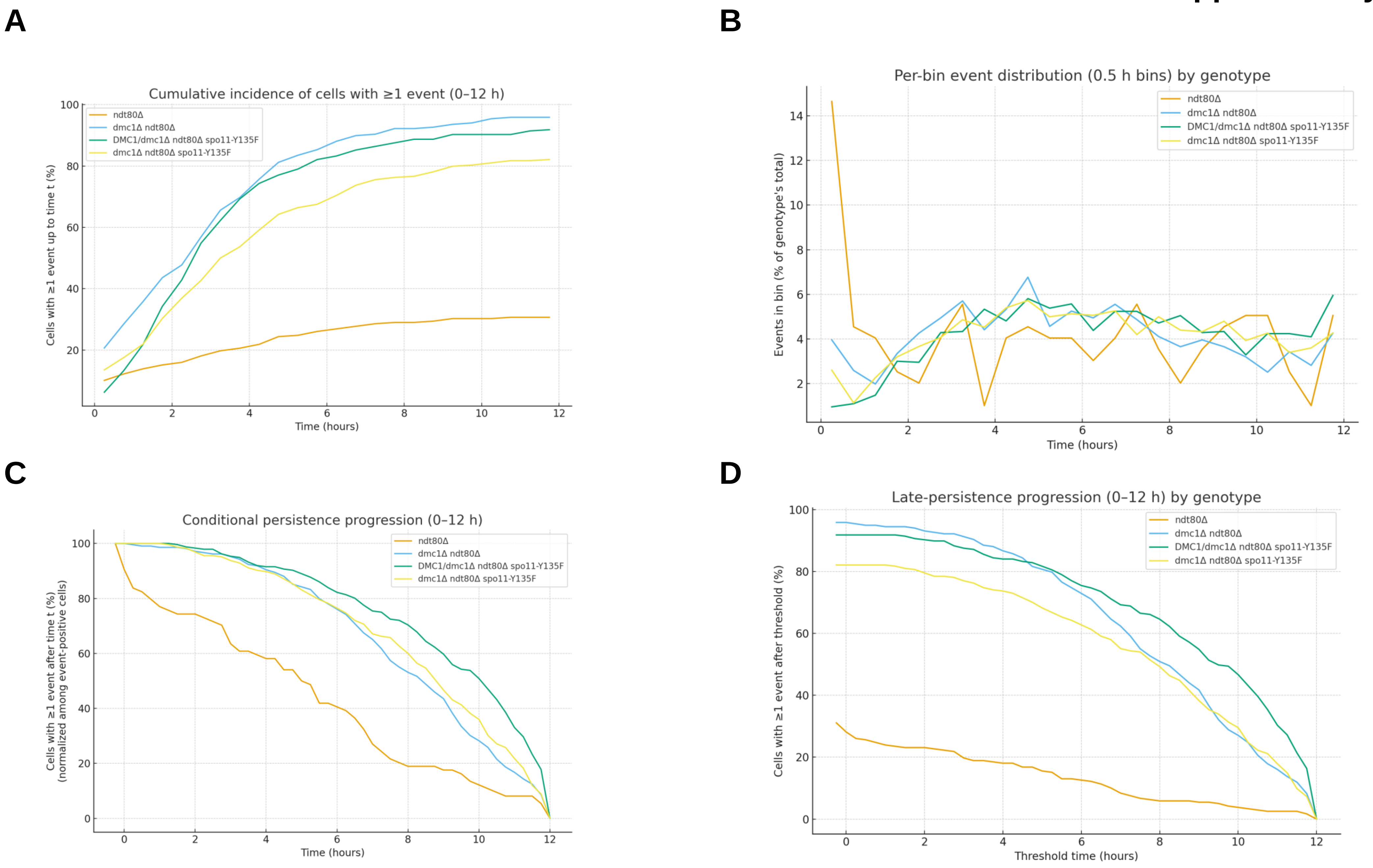

Table S1. Z-stack and time-lapse acquisition parameters

| Z-range (μm) | Section thickness (μm) | Number of sections | Number of positions | Number of time points |
|---|---|---|---|---|
| 5 | 0.35 | 15 | 10 | 49 |

Z-range, optical section thickness, number of Z-sections, positions per field and total number of time points used for all live-cell time-lapse acquisitions (15-min intervals over a 12-h window).

Supplementary Table S2A. Late-event fractions at 6.0 h and 9.0 h under Rec8–GFP prophase-I gating.

| Genotype | N cells | Cells with ≥1 event ≥6.0 h (n) | Fraction ≥6.0 h (%) | Cells with ≥1 event ≥9.0 h (n) | Fraction ≥9.0 h (%) |
|---|---|---|---|---|---|
| *ndt80Δ* | 238 | 30 | 12.6 | 14 | 5.9 |
| *dmc1Δ ndt80Δ* | 218 | 159 | 72.9 | 96 | 44.0 |
| *spo11-Y135F ndt80Δ* | 257 | 194 | 75.5 | 147 | 57.2 |
| *dmc1Δ spo11-Y135F ndt80Δ* | 274 | 172 | 62.8 | 114 | 41.6 |

$\chi^2$ tests comparing late-event fractions at 6.0 h and 9.0 h to the *ndt80Δ* reference.

Supplementary Table S2B.

| Comparison vs *ndt80Δ* | Threshold (h) | $\chi^2$ (df = 1) | p-value |
|---|---|---|---|
| *dmc1Δ ndt80Δ* | 6.0 | 170.6 | $5.34\times10^{-39}$ |
| *spo11-y135f ndt80Δ* | 6.0 | 197.2 | $8.46\times10^{-45}$ |
| *dmc1Δ spo11-y135f ndt80Δ* | 6.0 | 134.2 | $4.94\times10^{-31}$ |
| *dmc1Δ ndt80Δ* | 9.0 | 90.5 | $1.86\times10^{-21}$ |
| *spo11-y135f ndt80Δ* | 9.0 | 148.3 | $4.14\times10^{-34}$ |
| *dmc1Δ spo11-y135f ndt80Δ* | 9.0 | 86.7 | $1.27\times10^{-20}$ |

Late-event fractions at 6.0 h and 9.0 h were compared to the ndt80Δ reference using 2×2 $\chi^2$ tests (df = 1); all contrasts were highly significant ($p < 10^{-20}$).

Supplementary Table S3. Summary of last-event time (TLAST) per genotype under Rec8–GFP prophase-I gating.

| Genotype | n cells with ≥1 event | TLAST median (h) | TLAST IQR (h; Q1–Q3) | Mean TLAST (h) | SD TLAST (h) |
|---|---|---|---|---|---|
| *ndt80Δ* | 74 | 5.13 | 1.69–7.25 | 5.08 | 3.72 |
| *dmc1Δ ndt80Δ* | 209 | 8.50 | 6.25–10.50 | 8.12 | 2.79 |
| *spo11-Y135F ndt80Δ* | 236 | 10.25 | 7.44–11.50 | 9.16 | 2.83 |
| *spo11-y135f dmc1Δ ndt80Δ* | 225 | 9.00 | 6.25–11.00 | 8.37 | 2.90 |

TLAST was defined as the last frame (0–12 h, 15-min intervals) with a detectable nucleolar splitting event per cell. The table reports, for each genotype, the number of cells with ≥1 event, the median TLAST with interquartile range (IQR), and the mean ± SD.

Supplementary Table S4. Media

| Media | Components |
|---|---|
| YEPD | Yeast extract (1%), Bacto-peptone (2%), glucose (2%) |
| YEPD agar | Yeast extract (1%), Bacto-peptone (2%), glucose (2%), agar (2%) |
| YPA | Yeast extract (1%), Bacto-peptone (2%), Potassium acetate (1%) |
| SPM | Potassium acetate (1%) |
| Minimal medium | Yeast Nitrogen Base (-aa) 0.7%, glucose 2% and agar 2% |
